# ProAR: Probabilistic Autoregressive Modeling for Molecular Dynamics

**DOI:** 10.64898/2026.03.20.713063

**Authors:** Kaiwen Cheng, Yutian Liu, Zhiwei Nie, Mujie Lin, Yanzhen Hou, Yiheng Tao, Chang Liu, Jie Chen, Youdong Mao, Yonghong Tian

## Abstract

Understanding the structural dynamics of biomolecules is crucial for uncovering biological functions. As molecular dynamics (MD) simulation data becomes more available, deep generative models have been developed to synthesize realistic MD trajectories. However, existing methods produce fixed-length trajectories by jointly denoising high-dimensional spatiotemporal representations, which conflicts with MD’s frame-by-frame integration process and fails to capture time-dependent conformational diversity. Inspired by MD’s sequential nature, we introduce a new probabilistic autoregressive (ProAR) framework for trajectory generation. ProAR uses a dual-network system that models each frame as a multivariate Gaussian distribution and employs an anti-drifting sampling strategy to reduce cumulative errors. This approach captures conformational uncertainty and time-coupled structural changes while allowing flexible generation of trajectories of arbitrary length. Experiments on ATLAS, a large-scale protein MD dataset, demonstrate that for long trajectory generation, our model achieves a 7.5% reduction in reconstruction RMSE and an average 25.8% improvement in conformation change accuracy compared to previous state-of-the-art methods. For conformation sampling task, it performs comparably to specialized time-independent models, providing a flexible and dependable alternative to standard MD simulations.

## Introduction

Molecular dynamics (MD) simulations are commonly used to study the structural changes and movement patterns of biomolecules. By numerically integrating Newton’s equation of motion, MD simulations allow researchers to analyze protein conformations, drug-target interactions, and biological processes under equilibrium conditions (Alder and Wainwright 1959; Karplus and Kuriyan 2005; McCammon 1984). However, MD simulations face two main limitations: (1) the need for complex parameterization to accurately model non-covalent interactions, which restricts the size and complexity of systems that can be practically simulated; (2) significant temporal gaps, since many biologically important processes occur over timescales much longer than those accessible to standard MD techniques (Lewandowski et al. 2015; Kokkonen et al. 2018). Consequently, computing complete equilibrium trajectories remains computationally demanding and often unfeasible.

With advances in deep learning, models such as AlphaFold3 (Abramson et al. 2024) and RoseTTAFold All-Atom (Krishna et al. 2024) have achieved unprecedented accuracy in static biomolecular structure prediction. While they produce reliable structural snapshots, they cannot show how proteins physically morph between different functional states during biological processes (Nussinov et al. 2022). This limitation creates a significant gap in understanding functional mechanisms, as real proteins dynamically rearrange their structures to perform specific biological tasks.

Recent advances in deep learning have opened new pathways for exploring protein conformational landscapes. Early methods mainly focused on modifying AlphaFold’s inference protocol by strategically altering its multiple sequence alignment (MSA) inputs (Stein and Mchaourab 2022; Wayment-Steele et al. 2024). More recently, Distributional Graphformer (DiG) (Zheng et al. 2024) leverages a diffusion framework to iteratively refine a prior distribution toward the system’s equilibrium state, guided by molecular descriptors as conditional parameters. Meanwhile, the Str2Str framework (Lu et al. 2023) applies geometric perturbations to input structures, followed by denoising operations to navigate conformational landscapes.

Unlike methods that generate conformations through repeated i.i.d. sampling, we focus on directly generating MD trajectories. MDGEN (Jing et al. 2024) jointly tokenizes trajectories across spatial and temporal dimensions and uses stochastic interpolants (Ma et al. 2024) conditioned on specific keyframes to generate trajectories from Gaussian noise. Alphafolding (Cheng et al. 2025) utilizes motion guidance derived from historical observations to guide denoising during spatiotemporal (4D) diffusion. However, these denoising methods conflict with the sequential nature of traditional MD integration, resulting in two main limitations: (1) jointly denoising high-dimensional spatiotemporal representations adds substantial modeling complexity and fails to capture temporally correlated conformational diversity; and (2) training on fixed-length trajectories with a non-autoregressive design reduces inference flexibility, preventing the generation of variable-length trajectories.

At its core, conventional MD simulation can be formulated as a stochastic diffusion process (i.e., the Langevin equation (Schlick 2010; Allen and Tildesley 2017)), integrated sequentially frame by frame. Inspired by this, we suggest modeling MD trajectories in an autoregressive way. As shown in Figure 1, a simple solution would be to use a deterministic network (Li et al. 2025); however, this overlooks the inherent stochasticity of MD simulation and limits exploration of the free energy landscape. Instead, we introduce **Pro**babilistic **A**uto**R**egressive Modeling (**ProAR**), which explicitly models each frame as a multivariate Gaussian distribution from which samples are drawn. This probabilistic approach allows ProAR to capture conformational uncertainty and time-coupled structural changes, resulting in broader and more representative sampling of the energy landscape. Specifically, ProAR combines a dual-network system—a stochastic, time-conditioned interpolator and forecaster—with an anti-drifting sampling strategy. The interpolator predicts intermediate states as sparse-structured Gaussians by estimating their mean and covariance, while the forecaster infers future conformations through a corruption–refinement process conditioned on past observations. To prevent the accumulation of stochastic errors during autoregressive generation, the anti-drifting sampling strategy alternates between these two networks to perform intermediate frame interpolation and forward extrapolation. This enables flexible and stable generation of trajectories of arbitrary length.

**Figure 1.**
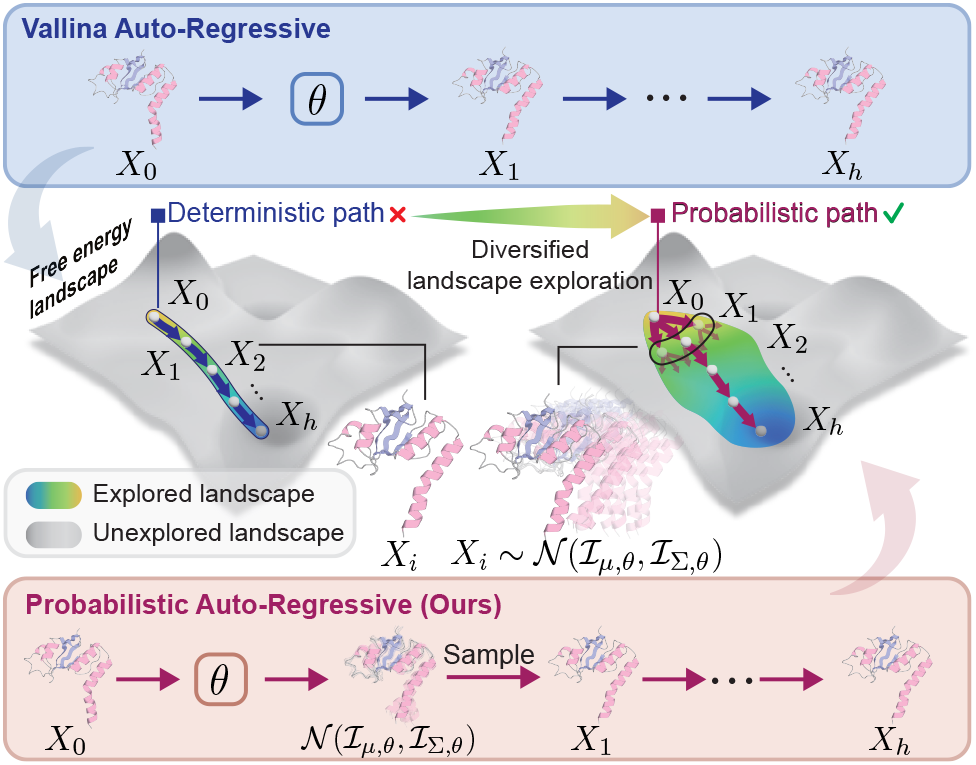
Comparison of autoregressive strategies for MD trajectory generation. Top: Vanilla AR produces a single deterministic path, limiting exploration of the free energy landscape. Bottom: Our probabilistic AR (ProAR) explicitly models each frame as a multivariate Gaussian, enabling sampling that captures conformational uncertainty and time-coupled structural changes, which in turn results in broader and more representative coverage of the landscape.

We evaluate ProAR on three representative protein dynamics tasks using the ATLAS dataset (Vander Meersche et al. 2024). First, in trajectory generation, ProAR surpasses MDGEN in both reconstruction fidelity and accuracy of conformation changes. Second, in conformation sampling, ProAR achieves performance comparable to specialized time-independent sampling models like AlphaFlow (Jing, Berger, and Jaakkola 2024), demonstrating its versatility beyond purely time-dependent tasks. Third, we leverage the interpolator to generate smooth transition pathways between distinct conformational states. In summary, ProAR introduces a novel probabilistic autoregressive paradigm that effectively captures time-dependent conformational variability and enables flexible, high-fidelity modeling of protein dynamics, serving as a practical and efficient alternative to MD simulation.

## Related Work

### Conformation Sampling

Molecular conformation sampling aims to generate time-independent ensembles. Early approaches such as MSA subsampling alter AlphaFold’s MSA input to produce diverse conformations through repeated inference (Stein and Mchaourab 2022; Wayment-Steele et al. 2024). Diffusion-based methods offer another route: Str2Str (Lu et al. 2023) applies denoising score matching, and CONFDIFF (Wang et al. 2024) adds MD energy-based guidance to bias sampling toward equilibrium. These approaches use limited dynamic cues from static structures and miss the richer patterns in MD simulations. With growing MD datasets, recent models train directly on trajectories. AlphaFlow (Jing, Berger, and Jaakkola 2024) and P2DFlow (Jin et al. 2025) improve diversity by adapting static predictors and fine-tuning them on MD data within a flow-matching framework. DiG (Zheng et al. 2024) regularizes the score to match MD force fields at small diffusion times and propagates this alignment through the Fokker-Planck equation. Although effective for equilibrium sampling, these models remain time-independent and cannot capture temporal correlations or transitions.

### Trajectory Generation

Instead of sampling conformations independently, recent methods focus on generating trajectories directly. Time-warp (Klein et al. 2023) uses normalizing flows within MCMC to propose large, transferable time jumps while preserving Boltzmann statistics. Several approaches are based on diffusion models, such as EquiJump (Costa et al. 2024), AlphaFolding (Cheng et al. 2025), and MDGEN (Jing et al. 2024). EquiJump employs SO(3)-equivariant stochastic interpolants for step-by-step trajectory generation, while AlphaFolding and MDGEN utilize spatiotemporal diffusion frameworks to model dynamics across space and time. AlphaFolding combines atomic grouping and motion alignment for temporally consistent generation, and MDGEN represents trajectories as joint spatiotemporal tokens with keyframe-conditioned stochastic interpolants. Though progress has been made, these approaches still encounter challenges with high complexity and insufficient temporal modeling due to joint denoising and fixed-length, non-autoregressive designs, which limit their flexibility and accuracy in capturing biomolecular dynamics. Recently, GST (Li et al. 2025) uses spatiotemporal Transformers with a Temporal Difference Graph to capture long-range dependencies; however, its deterministic autoregressive framework limits diversity. We introduce ProAR, which employs a dual-network design to model each frame as a multivariate Gaussian and adopts an anti-drifting sampling strategy to reduce cumulative errors, enabling diverse, time-coupled, and length-flexible trajectory generation.

## Method

### Problem Setup

Our objective is to learn the probability distribution of MD trajectories over a horizon of *H* time steps, denoted as *p*(**x**_1:*H*_ |**x**_0_, *c*), where **x**_0_ is the initial structure and *c* encodes conditioning information such as the biomolecular sequence. In this work, we primarily focus on protein structures. Following AlphaFold2 (Jumper et al. 2021), we parameterize the coordinates of heavy atoms by representing the backbone atoms (N–C*α*–C) of each residue as a local frame, constructed via a Gram–Schmidt process and referred to as a rigid frame. The global position and orientation of each rigid are described by a translation–rotation pair in SE(3) space, yielding a backbone conformation of a protein with *N* residues: **x** = (**T, R**) ∈ SE(3)^*N*^, where **T** ∈ℝ ^*N×*3^ and **R** ∈ SO(3)^*N*^ are the translation and rotation components. Full atomistic detail is further captured by up to seven torsional angles (*ϕ, ψ, ω, χ*_1_, …, *χ*_4_) per residue, describing bond rotations. Thus, the complete protein structure is parameterized as **x** = (**T, R**, *ϕ, ψ, ω, χ*_1_, …, *χ*_4_) ∈ (SE(3) × 𝕋^7*N*^. Existing diffusion models typically estimate the score functions of reverse-time marginals and model the joint distribution 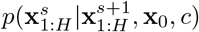, where *s* denotes the diffusion step, which requires simultaneous denoising across spatial and temporal dimensions. In contrast, to keep training tractable, we train over a shorter horizon *h* by learning *p*(**x**_*t*+1:*t*+*h*_ | **x**_*t*_, *c*) and apply the model autoregressively. This process begins with *p*(**x**_1:*h*_ | **x**_0_, *c*) and sequentially extends predictions until reaching the final horizon *H* during inference. For clarity, we omit *c* in later formulas, though it is always included as input.

### Probabilistic Autoregressive Modeling

Inspired by viewing MD simulations as stochastic diffusion processes, we introduce ProAR: a probabilistic autore-gressive framework with a dual-network design featuring a stochastic interpolator and forecaster. A dual-phase training approach enables ProAR to learn probabilistic distributions over conformations, while an anti-drifting sampling method supports stable and flexible generation of trajectories of arbitrary length.

#### Interpolator Training

We first train a time-conditioned stochastic interpolator ℐ_*ϕ*_ with parameter *ϕ* to interpolate between observed data snapshots, as shown in Figure 2a. Specifically, given a horizon *h*, the model is trained to ap-proximate intermediate states via 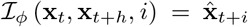 for *i* ∈ {1, …, *h* – 1} by optimizing the following objective:

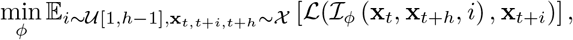

where **x**_*t,t*+*i,t*+*h*_ are samples drawn from the MD simulation dataset 𝒳, and ℒ denotes the loss function. A natural choice for ℒ is a deterministic structural loss, such as those proposed in AlphaFold2 (Jumper et al. 2021). However, this approach overlooks the intrinsic stochasticity of MD simulations, often resulting in mode collapse during training: the network degenerates to predicting only the ensemble mean, thereby losing the capacity to generate heterogeneous conformations. To address this, we enable ℐ_*ϕ*_ to explicitly model conformational uncertainty. Inspired by heteroscedastic regression (Gao, Barzel, and Yan 2024; Simpson, Vicente, and Campbell 2022), we represent each intermediate frame as a structured anisotropic multivariate Gaussian *p*(**x**_*t*+*i*_) = 𝒩 (***µ***_*t*+*i*_, **Σ**_*t*+*i*_), which captures time-dependent probabilistic motion patterns across different structural regions of the protein. We further decompose ℐ_*ϕ*_ into two branches: ℐ_***µ***,*ϕ*_ to predict the mean ***µ***_*t*+*i*_, and ℐ_Σ,*ϕ*_ to predict the covariance **Σ**_*t*+*i*_. Since the ground-truth covariance is unavailable, both branches are jointly supervised by minimizing the negative log-likelihood (NLL) of the predicted distribution (Nix and Weigend 1994):

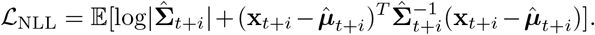

**Figure 2.**
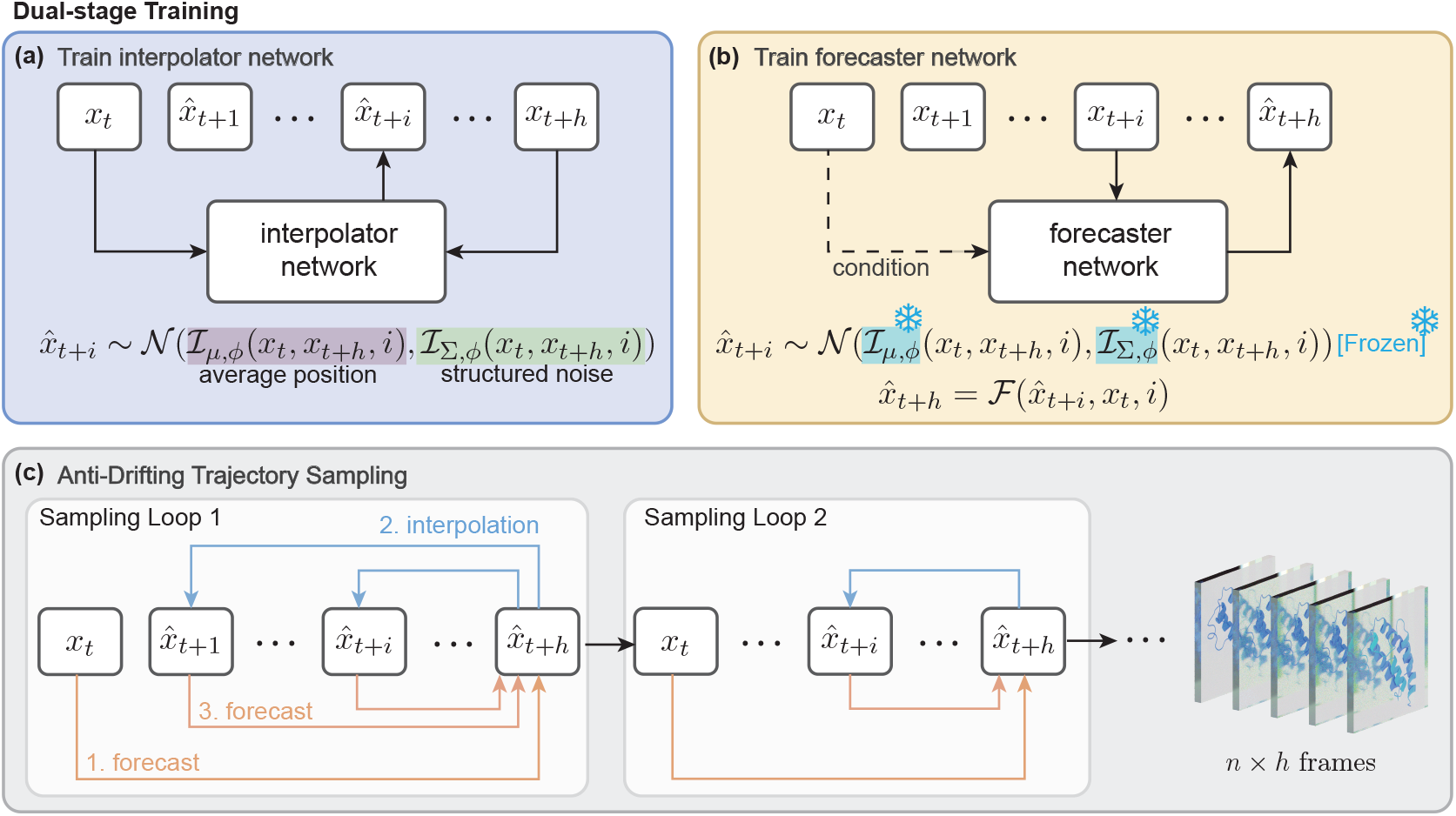
ProAR employs a dual-network system: an interpolator predicts intermediate frames **x**_*t*+*i*_ as structured multivariate Gaussians, estimating their mean and covariance from **x**_*t*_ and **x**_*t*+*h*_; a forecaster, conditioned on **x**_*t*_, takes the interpolator’s prediction 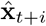 as input and produces a prediction for **x**_*t*+*h*_. During inference, alternating between the two networks allows for stable and flexible autoregressive generation, which we refer to as anti-drifting sampling.

In summary, the final objective combines the deterministic structural loss adapted from AlphaFold2 with the NLL term:

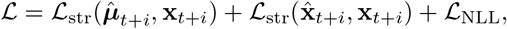

where 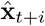 is sampled from 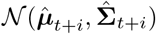 using the repa-rameterization trick and *ℒ*_str_ mainly consists of Frame aligned point error (FAPE), torsion angle loss, cross-entropy loss for distogram prediction, and violation loss. This design not only ensures that the interpolated states remain physically plausible and accurate, but also enables the model to capture structured noise and variability beyond deterministic averaged position.

More concretely, we model the residue-level probabilistic distribution where the mean ***µ***_*t*+*i*_ has size *N* and a full covariance **Σ**_*t*+*i*_ would need *N*^2^ parameters. To guarantee symmetry, positive definiteness, and efficient log-likelihood computation, we parameterize the covariance matrix by predicting its Cholesky factor **L**_Λ,*t*+*i*_ with 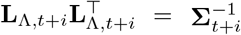. To further mitigate the quadratic complexity as *N* increases, we impose sparsity by only filling entries in **L**_Λ,*t*+*I*_ corresponding to residues within local neighborhoods, reflecting the localized correlations in protein dynamics. Further details for stabilizing training are described in the supplementary material.

#### Forecaster Training

In the second stage, we train a forecaster ℱ _*θ*_ with parameter *θ* to predict **x**_*t*+*h*_ such that 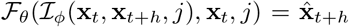 for *j*∈ {0, …, *h* − 1}, as shown in Figure 2b. Specifically, we seek to optimize the following objective with *ℐ*_*ϕ*_ frozen:

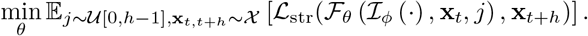

To incorporate boundary case, we define ℐ_*ϕ*_(**x**_*t*_,, 0):= **x**_*t*_. To enable the forecaster to identify the most probable future structure, we adopt a corruption–refinement paradigm inspired by prior independent sampling methods (Lu et al. 2023; Jin et al. 2025), but crucially make **x**_*t*_ and *j* explicit conditioning inputs. Specifically, we treat the interpolator output 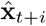 as the target for corruption, and generate a prior by adding Gaussian noise whose variance depends on *j*; smaller *j* (i.e., longer extrapolation horizons) are assigned larger variance to enhance conformational diversity. During refinement, the forecaster explicitly conditions on the historical structure **x**_*t*_ to ensure structural fidelity and guide predictions toward high-probability future states. Unlike independent conformation sampling methods that require multi-step denoising, our approach directly models time-coupled conformational changes by conditioning on both **x**_*t*_ and *j*, achieving accurate forecasting in a single forward pass.

#### Anti-Drifting Sampling

The stochasticity introduced by the interpolator and forecaster is crucial for probabilistic modeling. However, to ensure stable long-term generation, it is necessary to mitigate the risk of cumulative errors in the autoregressive process. To address this, we design an anti-drifting sampling strategy that alternately applies the interpolator and forecaster to perform intermediate frame interpolation and forward extrapolation, as shown in Figure 2c.

Starting from the initial condition **x**_0_, the forecaster ℱ_*θ*_ first predicts 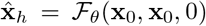. Given this forecast, the interpolator ℐ_*ϕ*_ produces the next intermediate frame 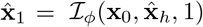. The prior forecast of **x**_*h*_ can now be refined with 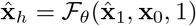. This alternating process continues until the interpolator predicts 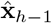 and the forecaster makes a final refinement for 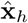. The process then repeats autoregressively, using 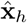 as the new starting point for subsequent loops. Notably, while ℱ_*θ*_ always forecasts the last frame of each loop, its prediction quality improves iteratively as the context moves closer in time to *t* + *h*, effectively reducing drift and maintaining temporal coherence throughout extended trajectories. To further prevent physically implausible structures—such as bond breaking or atomic clashes— during long trajectory generation, we follow the practice in AlphaFold2 by applying Amber relaxation at the end of each autoregressive loop using the AMBER99SB (Hornak et al. 2006) force field.

### Model Architecture

#### Shared Design

An overview of our model architecture is shown in Figure 3. Both the interpolator and forecaster share a common backbone built from SE(3)-equivariant blocks, combining Invariant Point Attention (IPA) (Abram-son et al. 2024) and E(n)-Equivariant Graph Neural Networks (EGNN) (Satorras, Hoogeboom, and Welling 2021). To enhance generalization, we initialize residue single and pair representations with ESM-2 language model embeddings (Lin et al. 2023), further enriched by encoded features from timestep, amino acid sequence, secondary structure type, and residue index—collectively providing temporal, sequential, and structural cues.

**Figure 3.**
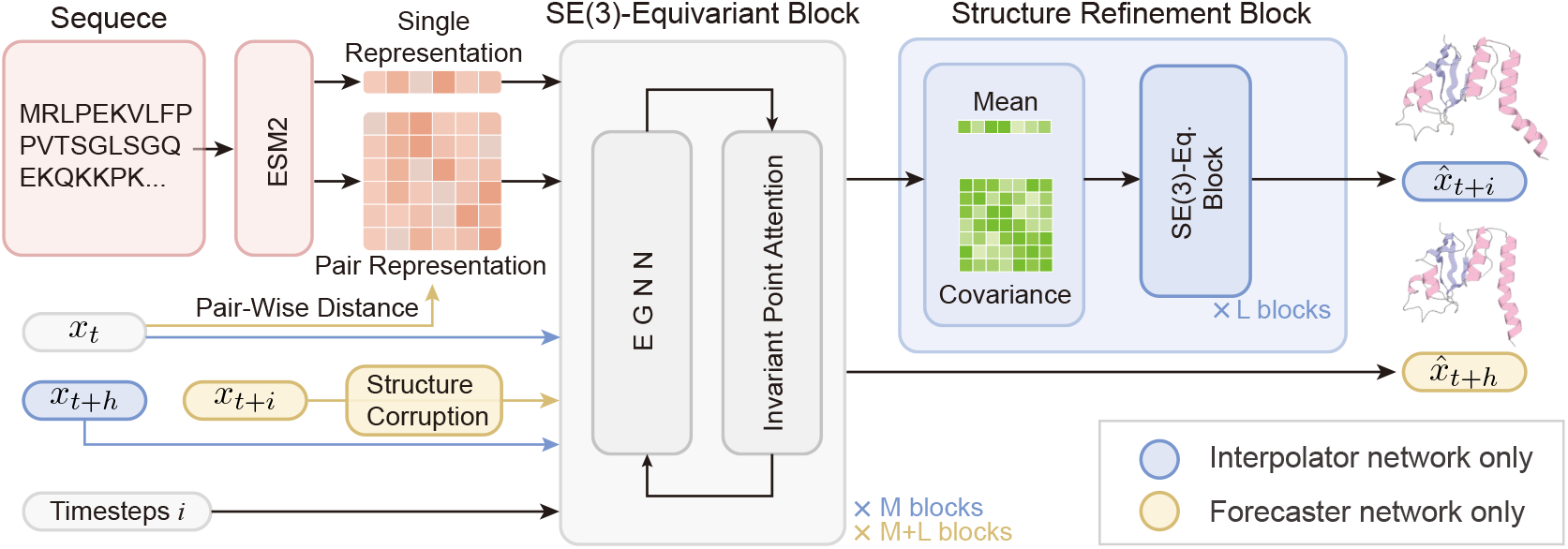
Architecture overview. Both networks utilize SE(3)-equivariant blocks that combine EGNN and IPA, with single and pair representations initialized from ESM-2 embeddings and encoded protein properties. In the interpolator, the first *M* blocks iteratively update **x**_*t*_ and **x**_*t*+*h*_, followed by *L* blocks that refine samples drawn from the predicted distribution. In the forecaster, corruption is applied via a stochastic perturbation kernel, and **x**_*t*_ is encoded through pairwise distances into the pair representation for conditioning.

#### Interpolator-Specific Design

In the interpolator, the SE(3)-eq. blocks simultaneously take both **x**_*t*_ and **x**_*t*+*h*_ as input. The EGNN update these two inputs in an alternating manner across successive blocks, while the IPA is modified to add a parallel point attention that allows it to fuse 3D context from both **x**_*t*_ and **x**_*t*+*h*_ into single representation, followed by two separate backbone updates for **x**_*t*_ and **x**_*t*+*h*_. After *M* SE(3)-eq. blocks, a learnable weighted average of 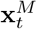and 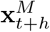 yields the distribution mean, while a Multi-Layer Perceptron (MLP) regresses the Cholesky factor of the covariance from the single representation Finally, *L* additional SE(3)-eq. blocks refine samples drawn from this distribution to produce the final violation-free predictions 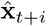.

#### Forecaster-Specific Design

Structural corruption is applied to the backbone conformation (**T, R**) ∈ SE(3)^*N*^ of 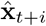 via a stochastic perturbation kernel that treats rotational and translational components independently. Specifically, **R** is perturbed element-wise using an isotropic Gaussian distribution on SO(3) (Yim et al. 2023), while **T** follows an Orn-stein–Uhlenbeck process (VP-SDE (Song et al. 2021)). The noise level linearly decreases as *j* increases. The corrupted conformation is then directly fed into (*M* + *L*) SE(3)-eq. blocks, while **x**_*t*_ is encoded through its C*α* pairwise distances and fused into the pair representation to provide guidance.

## Experiments

### Dataset

We evaluate our model using ATLAS (Vander Meersche et al. 2024), a large-scale molecular dynamics dataset comprising approximately 1300 proteins with diverse sizes and topologies. Each protein is represented by three independent 100 ns simulation trajectories. Following prior work (Jing, Berger, and Jaakkola 2024; Jing et al. 2024), we adopt the widely used dataset split by protein identity.

### Implementation Details

We set the horizon *h* = 6 with a 400 ps frame interval as a balanced choice: smaller *h* makes **x**_*t*_ and **x**_*t*+*h*_ too similar for meaningful interpolation, while larger *h* makes forecasting **x**_*t*+*h*_ from **x**_*t*_ harder. The interpolator is trained from scratch, with a stop-gradient technique (Stirn et al. 2023) to decouple mean and covariance branches for better covariance estimates. The forecaster is initialized from pretrained weights on the same dataset.

### Baselines

We benchmark ProAR against state-of-the-art deep learning methods for each task. For trajectory generation, we compare with MDGEN, a flow-based non-autoregressive model trained on ATLAS. AlphaFolding is excluded from comparison due to challenges in training and generalizing to larger proteins (*>*256 residues). For conformation sampling, we compare against AlphaFlow and CONFDIFF, which are flow- and diffusion-based, time-independent samplers fine-tuned on ATLAS. For conformation interpolation, since no existing baselines handle large proteins, we directly compare ProAR’s predictions with ground-truth MD trajectories.

### Trajectory Generation

We focus on generating long MD trajectories and assess model performance from two main aspects: (1) whether our model can accurately reproduce the ground truth MD trajectories; (2) whether it can precisely capture the extent of conformational changes.

#### Trajectory Reconstruction Fidelity

For each test protein, we generate a 250-frame trajectory from the initial structure of each of its three MD replicates, matching the 100 ns ATLAS reference trajectory. We use *R*_*s*_ to represent the average C*α*-RMSE (in angstroms, Å) over the first *s* steps. This benchmark evaluates how well the generated trajectories reproduce the ground truth MD simulations by measuring frame-wise deviation.

As shown in Table 1, ProAR outperforms MDGEN in reconstructing MD trajectories. Notably, our method excels at long-term prediction, lowering the *R*_250_ error from 3.813 to 3.529. While MDGEN generates all 250 frames simultaneously through an iterative denoising process, ProAR uses a multi-step autoregressive approach. These results demonstrate that, with the anti-drifting sampling strategy, ProAR effectively reduces cumulative error, leading to greater stability in long trajectory generation.

**Table 1:** Comparison of frame-wise C*α*-RMSE between MDGEN and ProAR, measured in angstroms (Å).

| | $R_{50}$ | $R_{100}$ | $R_{150}$ | $R_{200}$ | $R_{250}$ |
| --- | --- | --- | --- | --- | --- |
| MDGEN | 3.184 | 3.461 | 3.607 | 3.735 | 3.813 |
| ProAR | **3.120** | **3.292** | **3.380** | **3.472** | **3.529** |

#### Conformation Change Accuracy

Since both MD simulation and models produce stochastic trajectories, simply comparing frame-wise RMSE does not sufficiently determine whether the model accurately captures protein motion patterns in a probabilistic sense. To address this, we fit two separate PCA models per protein—one based on C*α* coordinates and another based on C*α* pairwise distances—using the triplicated ground truth MD trajectories. The generated trajectories were then projected into these PCA spaces. Conformation change accuracy was evaluated by first calculating: (1) Reference Displacement, which measures the *L*^2^-distance in PCA space from each frame to the initial frame; and (2) Stepwise Displacement, which measures the *L*^2^-distance between consecutive frames. Finally, the Hausdorff distance was used to compare the displacement distributions of the generated trajectories with those of the ground truth.

As shown in Table 2, the improvement indicates that ProAR more accurately captures probabilistic conformational uncertainty and time-coupled conformational change along the feature dimensions most relevant to the structural variance observed in MD. This result further demonstrates the advantage of our probabilistic modeling approach in providing richer and more accurate representations of dynamic molecular behavior. Figure 4 visualizes ensembles in generated trajectories, where ProAR captures broader conformational changes than MDGEN, reflecting key motions in both structured and disordered regions observed in the MD reference.

**Table 2:** Hausdorff distance measured in PCA space between generated and reference MD trajectories, based on Reference displacement (deviation from initial frame) and Step-wise displacement (deviation between consecutive frames).

| | PCA-2D($C\alpha$ pwd.) | | PCA-2D($C\alpha$ coord.) | |
| --- | --- | --- | --- | --- |
|  | Reference | Stepwise | Reference | Stepwise |
| MDGEN | 59.699 | 22.904 | 6.366 | 2.511 |
| ProAR | **40.607** | **18.722** | **4.467** | **1.926** |

**Figure 4.**
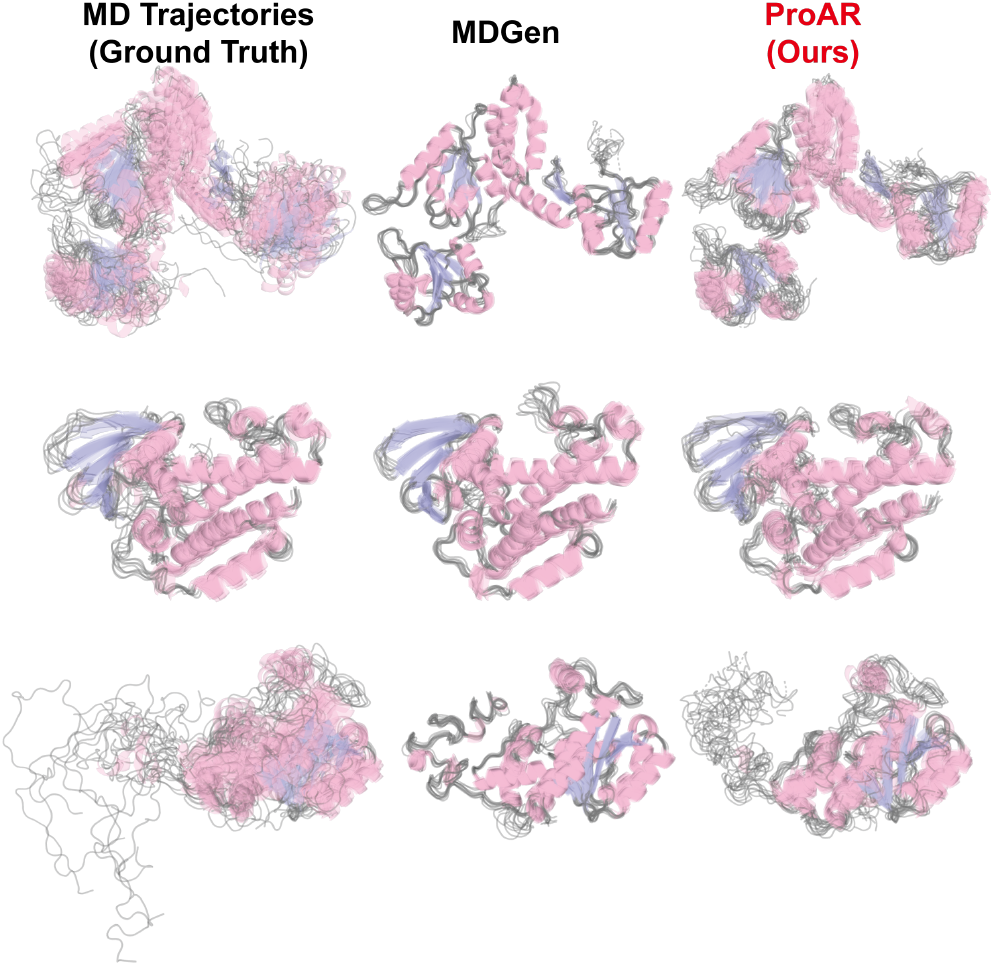
Visualization of three proteins from the ATLAS test set (6p5x-B, 7aex-A and 7c45-A). Each ensemble consists of 10 frames uniformly subsampled from the corresponding trajectory. MDGEN primarily exhibits minor fluctuations around the initial frame, whereas ProAR captures major movements in structured regions and better models the dynamic behavior of disordered regions.

### Conformation Sampling

We assess conformation sampling performance following the benchmark setup in (Jing, Berger, and Jaakkola 2024).

Specifically, for each protein, AlphaFlow and CONFD-IFF generate 250 independent samples, while MDGEN and ProAR produce a 250-frame trajectory starting from the initial frame of the MD simulation. Results are shown in Table 3. ProAR achieves performance comparable to state-of-the-art specialized models. Although AlphaFlow and CONFD-IFF are designed for time-independent conformation sampling and are additionally trained on Protein Data Bank (PDB) data, ProAR still attains the best results on 5 out of 7 metrics. This indicates that, although primarily designed for trajectory generation, ProAR effectively samples conformations that approximate the equilibrium distribution from MD simulations. Additionally, Figure 5 visualizes trajectories in PCA space, showing that ProAR explores a broader range of conformations, while MDGEN remains close to the initial structure and deviates from the true distribution.

**Table 3:**
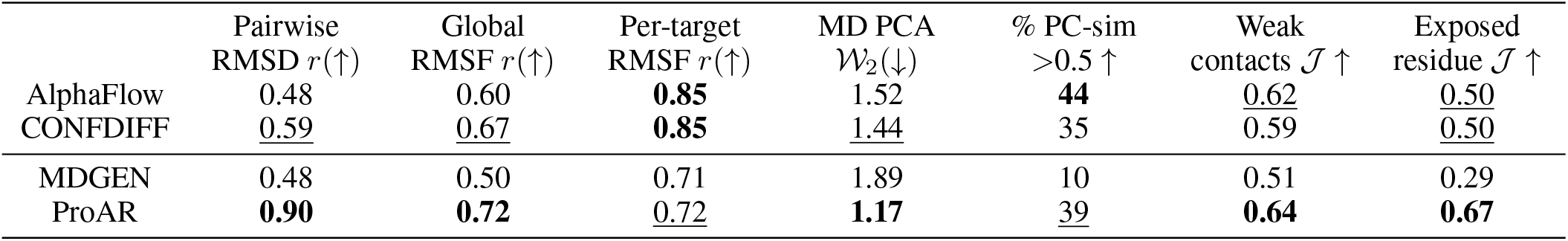
Conformation sampling benchmark. AlphaFlow and CONFDIFF are specialized time-independent SOTA models. The best scores are highlighted in **bold**, and the second-best scores are <u>underlined</u>.

**Figure 5.**
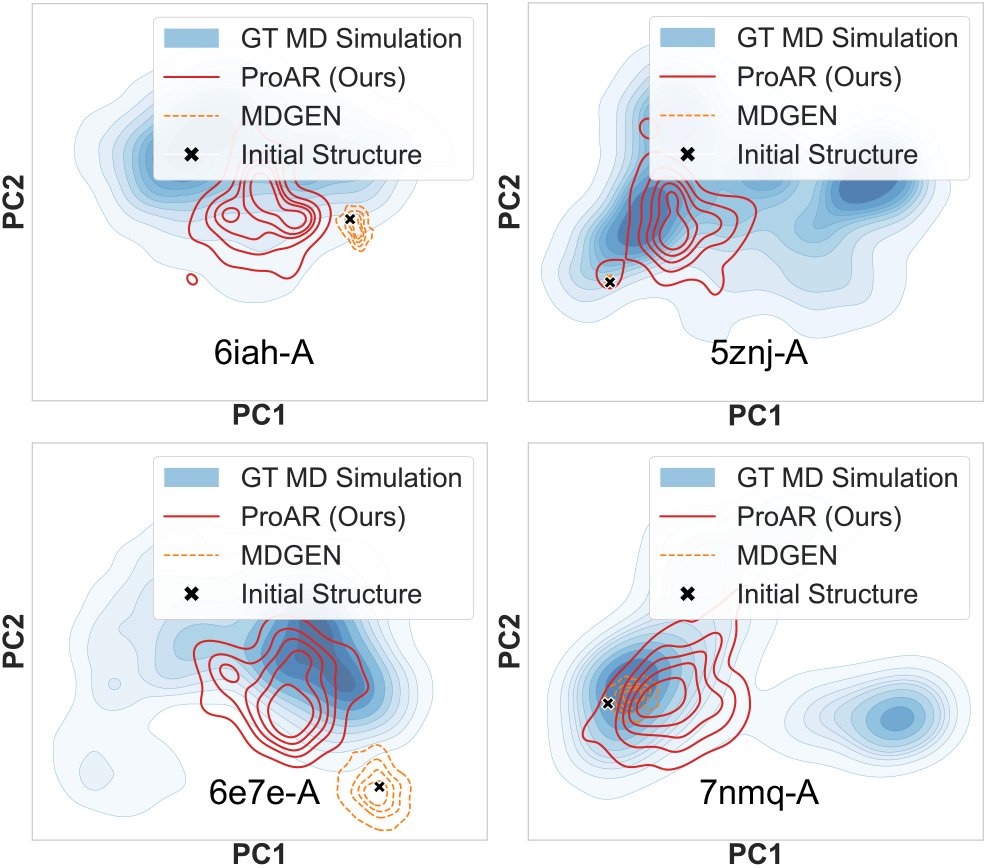
Representative trajectories illustrating the conformational space (in PCA-2D) explored by different models. Background contours indicate the density of the reference distribution obtained from MD simulations.

### Conformation Interpolation

In interpolation or transition path sampling, large free energy barriers make transitions rare and MD simulations expensive. We performed a preliminary study of ProAR’s interpolation ability. For each protein in the test set, we identified the frame with the largest RMSD relative to the first frame and considered these two frames as the start and end points. Using the interpolator, we then sampled a transition path connecting them. In these cases, the reference MD trajectories show significant conformational changes and clear state transitions. To evaluate whether our model produces smooth transitions toward the target end state, we measured the normalized *L*^2^ distance in PCA space between each intermediate frame and the start/end frames. As shown in Figure 6a, the distance to the start frame increases while the distance to the end frame decreases with frame index, indicating smooth and directed transitions. Figure 6b visualizes these intermediate pathways in PCA space, showing that ProAR’s transitions closely follow those observed in the reference MD trajectories.

**Figure 6.**
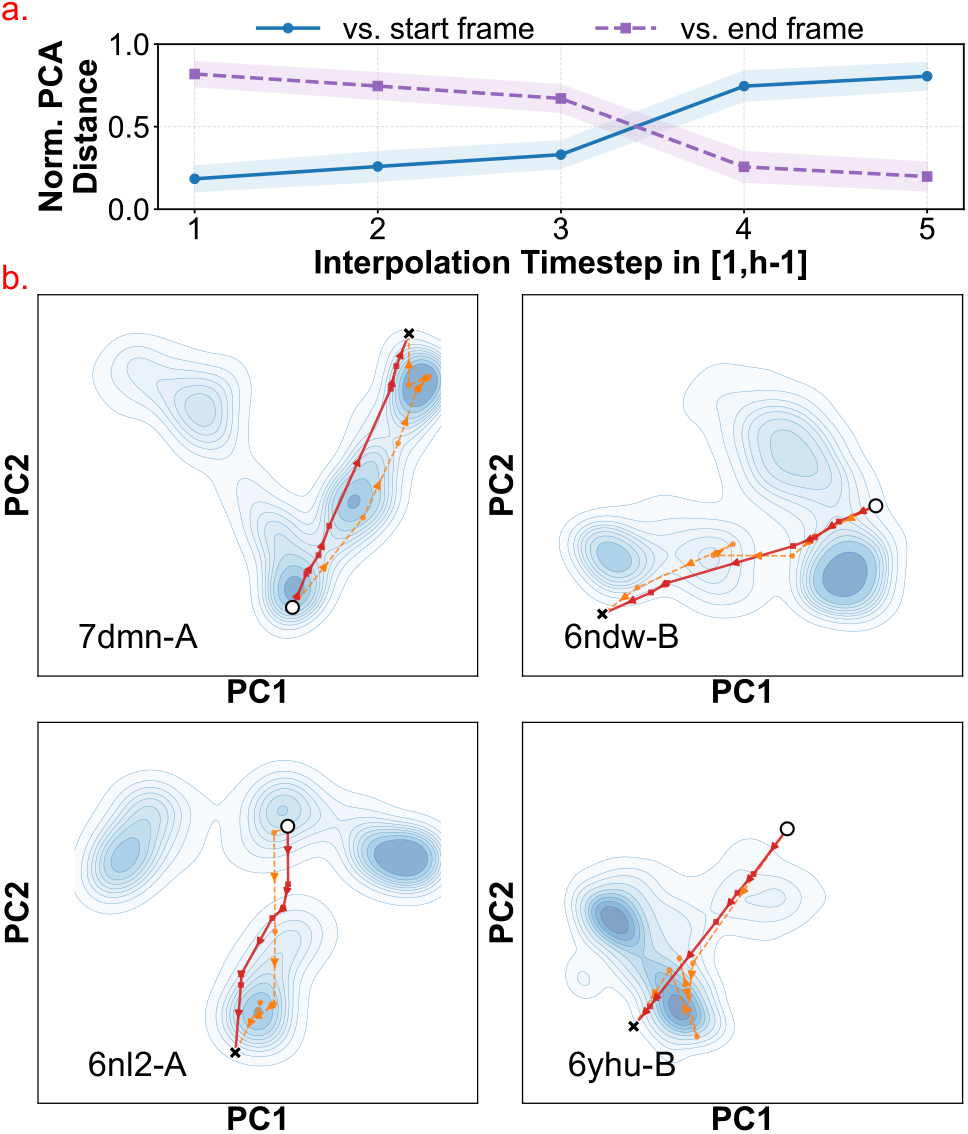
Conformation interpolation. (a) Normalized *L*^2^-distances in PCA space between intermediate frames and the start/end frames, averaged over 82 ATLAS test cases. (b) Example pathways showing that ProAR produces smooth transitions closely matching the dynamics in the MD reference.

## Conclusion

We introduce ProAR, a new probabilistic autoregressive framework for MD trajectory generation. ProAR uses a dual stochastic network structure for time-coupled probabilistic modeling, with an anti-drifting sampling strategy that ensures both flexibility and stability. Experiments show that ProAR: (1) surpasses SOTA methods in trajectory generation; (2) delivers competitive results in conformation sampling; and (3) effectively interpolates between conformations. ProAR highlights the importance of probabilistic autoregressive modeling in molecular simulation, providing an efficient and adaptable approach to capturing dynamics.

## Acknowledgments

This work was supported in part by Shenzhen KQTD (No.20240729102051063), Shenzhen Medical Research Funds of China (No. B2302037), National Natural Science Foundation of China (grant number 12125401), Natural Science Foundation of China (No. 61972217, 32071459, 62176249, 62006133, 62271465, 62406167), National Key Research and Development Program of China (grant number 2023YFF1204400 and 2023YFF1204401), and AI for Science (AI4S)-Preferred Program, Peking University Shenzhen Graduate School, China.

