## Supplementary Material for "ProAR: Probabilistic Autoregressive Modeling for Molecular Dynamics"

### Supplementary Material of
### ProAR: Probabilistic Autoregressive Modeling for Molecular Dynamics

**Kaiwen Cheng**[1,8*], **Yutian Liu**[2*], **Zhiwei Nie**[1,8*], **Mujie Lin**[1,8], **Yanzhen Hou**[3], **Yiheng Tao**[2], **Chang Liu**[7†], **Jie Chen**[1,8†], **Youdong Mao**[3,4,5,6,8†], **Yonghong Tian**[1,2,8†]

[1]School of Electronic and Computer Engineering, Peking University, Shenzhen, China
[2]School of Computer Science, Peking University, Beijing, China
[3]School of Physics, Peking University, Beijing, China
[4]Peking-Tsinghua Joint Center for Life Sciences, Peking University, Beijing, China
[5]Center for Quantitative Biology, Peking University, Beijing, China
[6]National Biomedical Imaging Center, Peking University, Beijing, China
[7]Department of Automation and BNRist, Tsinghua University, Beijing, China
[8]AI for Science (AI4S)-Preferred Program, Peking University Shenzhen Graduate School, China
{kaiwencheng, yutianliu}@stu.pku.edu.cn, {zhiweiNie, jiechen2019, ymao, yhtian}@pku.edu.cn,


#### Method Details

##### Detailed Model Architectures

The following three sections detail the core design motivations and architectures of the encoding layer, the interpolator, and the forecaster. Table 1 provides the hyperparameter configurations of the interpolator and the forecaster.

**Encoding Layer.** The protein-specific single and pair representations are first derived from ESM-2 (Lin et al. 2023): the single representation comes from the sequence embeddings in the representation layer, while the pair representation is initialized using the model's unsupervised self-attention maps. To further enrich these representations, we incorporate temporal, sequential, and structural cues. Specifically, temporal cues are introduced through sinusoidal timestep embeddings; sequential cues include sinusoidal positional embeddings of residue indices combined with learnable embeddings for the 20 standard amino acid types; and structural cues are captured by learnable embeddings corresponding to secondary structure types (helix, strand, or coil), computed using DSSP (Kabsch and Sander 1983). These additional features are concatenated with the single representation. To capture detailed pairwise interactions between residues, we compute the pairwise distances based on the C$\alpha$ coordinates of $\mathbf{x}_t$. These distances are then discretized into bins and transformed using a radial basis function (RBF). The resulting C$\alpha$ pairwise features are concatenated with the pair representation, enriching it with coarse-grained spatial context.

**Interpolator.** The interpolator jointly updates $\mathbf{x}_t$ and $\mathbf{x}_{t+h}$ within the first $M$ SE(3)-equivariant blocks. Specifically, the EGNN alternate their updates between these two inputs across successive blocks. The IPA is modified to add a parallel point attention, enabling it to fuse 3D context from both $\mathbf{x}_t$ and $\mathbf{x}_{t+h}$ into the single representation, as detailed in Algorithm 1. $\{\mathbf{s}_i\}$ and $\{\mathbf{z}_{ij}\}$ denote the single and pair representation. $\{T_{\mathbf{x}_t,i}\}$ and $\{T_{\mathbf{x}_{t+h},i}\}$ represent the rigid frames of $\mathbf{x}_t$ and $\mathbf{x}_{t+h}$, respectively. The settings $N_{\text{head}} = 12, c = 16, N_{\text{query points}} = 4$, and $N_{\text{point values}} = 8$ are kept consistent with the original AlphaFold2 design. Additionally, following Diffusion Transformer (Peebles and Xie 2023), we replace the standard layer norm in the original IPA with zero-initialized adaptive layer norm (adaLN-Zero) block to better incorporate timestep information.

After the updates through $M$ SE(3)-equivariant blocks, two linear layers independently regress per-residue average weights from $\{\mathbf{s}_i\}^M$, which are then used to perform translation averaging and rotation averaging for $\mathbf{x}_t^M$ and $\mathbf{x}_{t+h}^M$, yielding the predicted mean of the distribution.

For modeling the covariance of the distribution, we parameterize it via the Cholesky decomposition of the precision matrix (Simpson, Vicente, and Campbell 2022; Dorta et al. 2018; Williams 1996), i.e., the network directly predicts $\mathbf{L}_{\Lambda,t+i}$, where $\mathbf{L}_{\Lambda,t+i}\mathbf{L}_{\Lambda,t+i}^\top = \mathbf{\Sigma}_{t+i}^{-1}$ and $\mathbf{L}_{\Lambda,t+i}$ is a lower triangular matrix. Directly predicting the Cholesky factor ensures that the resulting precision matrix is symmetric, while enforcing positive diagonal elements guarantees positive definiteness. This parameterization facilitates straightforward computation of the log-likelihood of the predicted Gaussian. However, despite the Cholesky factor being lower triangular, its number of elements still scales quadratically with the number of residues $N$, making direct estimation impractical for large proteins. To address this, for each residue, we construct a sparse Cholesky matrix by filling only the entries corresponding to a small local neighborhood along the protein sequence, while preserving the lower-

*These authors contributed equally.
†Corresponding authors.

Copyright © 2026, Association for the Advancement of Artificial Intelligence (www.aaai.org). All rights reserved.

Algorithm 1: Modified IPA with two parallel point attention branches.

**Require:** $\{\mathbf{s}_i\}, \{\mathbf{z}_{ij}\}, \{T_{\mathbf{x}_t,i}\}, \{T_{\mathbf{x}_{t+h},i}\}, N_{\text{head}}, c, N_{\text{query points}}, N_{\text{point values}}$

$\mathbf{q}_i^h, \mathbf{k}_i^h, \mathbf{v}_i^h = \text{LinearNoBias}(\mathbf{s}_i)$

$\vec{\mathbf{q}}_{\mathbf{x}_t,i}^{hp}, \vec{\mathbf{k}}_{\mathbf{x}_t,i}^{hp} = \text{LinearNoBias}(\mathbf{s}_i)$

$\vec{\mathbf{v}}_{\mathbf{x}_t,i}^{hp} = \text{LinearNoBias}(\mathbf{s}_i)$

$\vec{\mathbf{q}}_{\mathbf{x}_{t+h},i}^{hp}, \vec{\mathbf{k}}_{\mathbf{x}_{t+h},i}^{hp} = \text{LinearNoBias}(\mathbf{s}_i)$

$\vec{\mathbf{v}}_{\mathbf{x}_{t+h},i}^{hp} = \text{LinearNoBias}(\mathbf{s}_i)$

$b_{ij}^h = \text{LinearNoBias}(\mathbf{s}_i)$

$w_C = \sqrt{\frac{2}{9N_{\text{query points}}}}, w_L = \sqrt{\frac{1}{3}}$

$a_{ij}^h = \text{softmax}_j(w_L(\frac{1}{\sqrt{c}}\mathbf{q}_i^{h\top}\mathbf{k}_j^h + b_{ij}^h$
$-\frac{\gamma^h w_C}{4}\sum_p \left\|T_{\mathbf{x}_t,i} \circ \vec{\mathbf{q}}_{\mathbf{x}_t,i}^{hp} - T_{\mathbf{x}_t,j} \circ \vec{\mathbf{k}}_{\mathbf{x}_t,j}^{hp}\right\|^2$
$-\frac{\gamma^h w_C}{4}\sum_p \left\|T_{\mathbf{x}_{t+h},i} \circ \vec{\mathbf{q}}_{\mathbf{x}_{t+h},i}^{hp} - T_{\mathbf{x}_{t+h},j} \circ \vec{\mathbf{k}}_{\mathbf{x}_{t+h},j}^{hp}\right\|^2))$

$\tilde{\mathbf{o}}_i^h = \sum_j a_{ij}^h \mathbf{z}_{ij}$

$\mathbf{o}_{\mathbf{x}_t,i}^h = \sum_j a_{ij}^h \mathbf{v}_{\mathbf{x}_t,j}^h$

$\mathbf{o}_{\mathbf{x}_{t+h},i}^h = \sum_j a_{ij}^h \mathbf{v}_{\mathbf{x}_{t+h},j}^h$

$\vec{\mathbf{o}}_{\mathbf{x}_t,i}^{hp} = T_{\mathbf{x}_t,i}^{-1} \circ \sum_j a_{ij}^h \left(T_{\mathbf{x}_t,j} \circ \vec{\mathbf{v}}_{\mathbf{x}_t,j}^{hp}\right)$

$\vec{\mathbf{o}}_{\mathbf{x}_{t+h},i}^{hp} = T_{\mathbf{x}_{t+h},i}^{-1} \circ \sum_j a_{ij}^h \left(T_{\mathbf{x}_{t+h},j} \circ \vec{\mathbf{v}}_{\mathbf{x}_{t+h},j}^{hp}\right)$

$\tilde{\mathbf{s}}_i = \text{Linear}(\text{concat}_{h,p}(\tilde{\mathbf{o}}_i^h, \mathbf{o}_{\mathbf{x}_t,i}^h, \vec{\mathbf{o}}_{\mathbf{x}_t,i}^{hp}, \left\|\vec{\mathbf{o}}_{\mathbf{x}_t,i}^{hp}\right\|$
$\mathbf{o}_{\mathbf{x}_{t+h},i}^h, \vec{\mathbf{o}}_{\mathbf{x}_{t+h},i}^{hp}, \left\|\vec{\mathbf{o}}_{\mathbf{x}_{t+h},i}^{hp}\right\|))$

**return** $\{\tilde{\mathbf{s}}_i\}$

triangular structure. This sparse representation allows us to compactly parameterize the matrix by predicting only the non-zero elements: specifically, for each residue $i$, we use $\mathbf{s}_i^M$ to predict its diagonal entry and the off-diagonal terms representing interactions with its four sequential neighbors.

While sparsification facilitates efficient computation, limited interactions is insufficient to capture the conformational heterogeneity shaped by complex protein topologies. As a result, the learned noise may violate basic structural and density constraints commonly observed in realistic protein conformations, leading to physically implausible outcomes such as bond breakage, atomic clashes, or violations from expected size scaling laws $R_g \propto N^\nu$ with $\nu \approx 0.4$ (Hong and Lei 2009; Tanner 2016), where $R_g$ stands for radius of gyration. To this end, we adopt the $R_g$-confined globular covariance model proposed in Chroma (Ingraham et al. 2023) as a prior for our learnable covariance. This prior (1) is SO(3)-invariant, (2) enforces protein chain connectivity and radius of gyration statistics, and (3) can be computed in linear time. By incorporating such structural prior, our model avoids allocating capacity and training time to re-learn well-established geometric constraints, allowing it instead to focus on capturing conformation uncertainty and time-coupled structural changes.

**Forecaster.** In the structure corruption process, we introduce higher noise level specifically to the coil regions of the protein, encouraging the forecaster to better capture their inherently flexible and disordered dynamics. The architecture mirrors that of P2DFlow (Jin et al. 2025): it comprises $M + L$ SE(3)-equivariant blocks that process $\mathbf{x}_{t+i}$, guided by the pairwise distance features computed from $\mathbf{x}_t$. In contrast to the interpolator which performs iterative backbone updates at each block, the forecaster applies a single update only at the final block. Following (Jin et al. 2025), we project each protein's MD simulation ensemble onto a 2D space defined by the $R_g$ and the RMSD from the crystal structure. We then estimate the Gaussian kernel density over this 2D map and apply the Boltzmann equation to obtain "approximate energy" values. During training, the approximate energy of $\mathbf{x}_{t+h}$ is used as an additional input: after each IPA block, an energy adapter performs cross-attention between this energy and the single representation. At inference time, the energy is sampled once from the Boltzmann distribution and kept fixed throughout each autoregressive loop.

#### Training and Sampling Details.

**Overall Training Configurations.** We train on the ATLAS dataset (Vander Meersche et al. 2024), following the train–validation–test split used in previous works (Jing, Berger, and Jaakkola 2024; Jing et al. 2024; Jin et al. 2025; Wang et al. 2024). The interpolator is trained from scratch, while the forecaster is initialized with pretrained weights from P2DFlow (Jin et al. 2025). Each MD trajectory in the training set is segmented into overlapping samples of length $h + 1$ with a stride of 400 ps, consistent with the setting in MDGEN (Jing et al. 2024).

During training, we adopt a variable batch size strategy by dynamically packing different proteins with similar residue counts into a single minibatch, while keeping the total number of residues below a predefined threshold: 1000 for the interpolator and 800 for the forecaster. We use the AdamW optimizer with weight decay $1 \times 10^{-4}$ and betas $[0.9, 0.99]$. The learning rate follows a cosine decay schedule with 200 warm-up steps, with a minimum of $5 \times 10^{-4}$ and a maximum of $1 \times 10^{-3}$. All training and sampling were carried out using 8 NVIDIA A800 GPUs. The following three sections detail the training process for the interpolator and forecaster, as well as the implementation of the anti-drifting sampling strategy.

**Interpolator Training.** For clarity, we adopt the following notation: let $f_{\text{trunk}}$ denote the shared representation learner—including the encoding layer and $M$ SE(3)-equivariant blocks. The mean parameter head is denoted as $f_{\boldsymbol{\mu}}$, and the covariance parameter head as $f_{\boldsymbol{\Sigma}}$. Together, $f_{\text{trunk}}$ and $f_{\boldsymbol{\mu}}$ constitute the mean branch $\mathcal{I}_{\boldsymbol{\mu},\phi}$, while $f_{\text{trunk}}$ and $f_{\boldsymbol{\Sigma}}$ form the covariance branch $\mathcal{I}_{\boldsymbol{\Sigma},\phi}$. A naive approach would train both heads end-to-end under the same negative log-likelihood objective. However, this approach allows the network to compensate for poor mean predictions by inflating the predicted variance, thus maintaining a low loss without truly improving the mean accuracy. Prior studies (Seitzer et al. 2022; Stirn et al. 2023) show that such coupling of-

| Hyperparameters | Interpolator | Forecaster |
|---|---|---|
| Number of SE(3)-equivariant blocks | $M = 4, L = 2$ | 6 |
| Single representation channel dimension | 384 | 256 |
| Pair representation channel dimension | 128 | 128 |
| Number of pairwise distance bins for $\mathbf{x}_t$ | 22 | 64 |
| IPA hidden channel dimension | 16 | 128 |
| Number of IPA heads | 12 | 16 |
| Number of IPA query points | 4 | 32 |
| Number of IPA value points | 8 | 8 |
| Number of Transformer layers | 2 | 1 |
| Number of Transformer attention heads | 4 | 16 |
| Number of EGNN layers | 2 | 2 |
| Dropout rate | 0.2 | 0.2 |

Table 1: Hyperparameter choices. $M$ and $L$ denote the number of SE(3)-equivariant backbone blocks and refinement blocks, respectively. For the interpolator, the Transformer applies self-attention to the single representation after each IPA block. For the forecaster, it refers to the energy adapter that performs cross-attention between the single representation and the approximate energy.

ten degrades the fidelity of mean estimates. To address this, (Stirn et al. 2023) propose decoupling the mean and variance heads by introducing a stop-gradient operation between the variance head and the shared trunk. This allows the variance head to benefit from the trunk's learned features, while preventing it from biasing the mean predictions. We adopt this strategy in the interpolator. Accordingly, our negative log-likelihood (NLL) loss term can be reformulated as:

$$\mathcal{L}_{\text{NLL}} = \mathbb{E}[-\log \mathcal{N}(\mathbf{x}_{t+i}; \lfloor f_{\boldsymbol{\mu}} \left(f_{\text{trunk}} \left(\mathbf{x}_t, \mathbf{x}_{t+i}, i\right)\right) \rfloor, \\ f_{\boldsymbol{\Sigma}} \left(\lfloor f_{\text{trunk}} \left(\mathbf{x}_t, \mathbf{x}_{t+i}, i\right) \rfloor\right))],$$

where $\lfloor \cdot \rfloor$ denotes the stop gradient operation. In practice, we first pretrain the mean branch $\mathcal{I}_{\boldsymbol{\mu},\phi}$ from scratch using only a structural loss $\mathcal{L}_{\text{str}}(\hat{\boldsymbol{\mu}}_{t+i}, \mathbf{x}_{t+i})$. Once the mean branch approaches convergence, we introduce the covariance branch $\mathcal{I}_{\boldsymbol{\Sigma},\phi}$ and add the other two loss terms, $\mathcal{L}_{\text{str}}(\hat{\mathbf{x}}_{t+i}, \mathbf{x}_{t+i})$ and $\mathcal{L}_{\text{NLL}}$.

**Forecaster Training.** For the forecaster, we initialize with pretrained weights from P2DFlow (Jin et al. 2025), which was trained on the same dataset split used in this work. P2DFlow is a generative model based on SE(3) flow matching, originally designed for time-independent conformation sampling. In the original formulation, the timestep is treated as a continuous variable within the flow matching framework. In our adaptation, we reinterpret this timestep $j$ as a discrete index representing different extrapolation horizons. Through continued training under this new formulation, the pretrained model is extended from purely time-independent sampling to time-dependent trajectory forecasting—predicting the most probable future conformations conditioned on historical observation $\mathbf{x}_t$ and the timestep $j$.

**Anti-Drifting Sampling.** The detailed anti-drifting sampling process is presented in Algorithm 2. Intuitively, the forecaster iteratively improves its prediction of $\mathbf{x}_{t+h}$ as the process comes closer in time to $t + h$. This motivates line 9 of Algorithm 2, where the final forecast of $\mathbf{x}_{t+h}$ can be used to fine-tune intermediate predictions.

Algorithm 2: Anti-drifting sampling.

**Require:** Initial structure $\hat{\mathbf{x}}_0 := \mathbf{x}_0$, training and inference horizon $h$ and $H$.
1: # Autoregressive loop:
2: **for** $t = 0, h, 2h, \ldots, (\lceil H/h \rceil - 1)h$ **do**
3: # Sampling loop for time steps $t + 1, \ldots, t + h$:
4: **for** $j = 0, 1, \ldots, h - 1$ **do**
5: $\hat{\mathbf{x}}_{t+h} \leftarrow \mathcal{F}_\theta(\hat{\mathbf{x}}_{t+j}, \hat{\mathbf{x}}_t, j)$ # (Refine) forecast
6: $\hat{\mathbf{x}}_{t+j+1} = \mathcal{I}_\phi \left(\hat{\mathbf{x}}_t, \hat{\mathbf{x}}_{t+h}, j + 1\right)$ # Interpolate
7: **end for**
8: # Optional refinement
9: $\hat{\mathbf{x}}_{t+k} \leftarrow \mathcal{I}_\phi \left(\hat{\mathbf{x}}_t, \hat{\mathbf{x}}_{t+h}, k\right), \forall k \in \{1, \ldots, h - 1\}$
10: # Perform Amber Relaxation
11: $\hat{\mathbf{x}}_{t+h} \leftarrow \text{Relax}(\hat{\mathbf{x}}_{t+h})$
12: **end for**
13: **return** $\hat{\mathbf{x}}_{1:H}$

#### Metrics

**Frame-wise RMSE.** We evaluate trajectory reconstruction fidelity using a frame-wise RMSE metric, computed by averaging over all frames, repeats, and proteins in the test set, as defined in the following equation:

$$\text{RMSE} = \frac{1}{M} \sum_{(p,r,t)} \sqrt{\frac{1}{N} \sum_{i=1}^{N} \left\| \hat{\mathbf{x}}_{\mathtt{C}\alpha,i}^{(p,r,t)} - \mathbf{x}_{\mathtt{C}\alpha,i}^{(p,r,t)} \right\|^2},$$

where $M$ sums over all frames indexed by protein $p$, repeat $r$, and timestep $t$. $N$ is the number of $\mathtt{C}\alpha$ atoms in protein $p$. $\hat{\mathbf{x}}_{\mathtt{C}\alpha,i}^{(p,r,t)}$ and $\mathbf{x}_{\mathtt{C}\alpha,i}^{(p,r,t)}$ are the predicted and ground-truth coordinates of $\mathtt{C}\alpha$ atom $i$.

**PCA Projection.** We fit two PCA-2D models per protein—one on $\mathtt{C}\alpha$ coordinates and another on $\mathtt{C}\alpha$ pairwise distances—using the triplicate ground-truth MD trajectories from ATLAS. All conformations are first aligned to the reference structure (i.e., the input structure used to initiate the simulation) before computing the PCA. Sampled conformations are likewise aligned prior to PCA projection. Performing analyses in the PCA space allows us to focus on the principal dimensions that best capture the structural variations observed in MD simulations.

**Hausdorff Distance.** The Hausdorff distance is used to quantify the similarity between the displacement sets of the generated trajectories and the ground truth. Specifically, given two point sets $\mathcal{A} = \{a_1, \dots, a_p\}$ and $\mathcal{B} = \{b_1, \dots, b_q\}$, the Hausdorff distance is defined as:

$$\mathrm{H}(\mathcal{A}, \mathcal{B}) = \max\big(\mathrm{h}(\mathcal{A}, \mathcal{B}),\ \mathrm{h}(\mathcal{B}, \mathcal{A})\big),$$

where $\mathrm{h}(\mathcal{A}, \mathcal{B}) = \max_{a \in \mathcal{A}} \min_{b \in \mathcal{B}} \|a - b\|, \mathrm{h}(\mathcal{B}, \mathcal{A}) = \max_{b \in \mathcal{B}} \min_{a \in \mathcal{A}} \|b - a\|$. Here, $\|\cdot\|$ denotes the Euclidean distance. The Hausdorff distance measures the maximum deviation between two point sets; a smaller value indicates that the generated trajectories more faithfully capture the true conformation changes.

#### Additional Visualization Results

**Additional PCA Visualization.** We additionally provide randomly selected PCA plots in Figure 1.

**Additional Interpolation Result.** We include additional visualization on interpolation results, as shown in Figure 2.

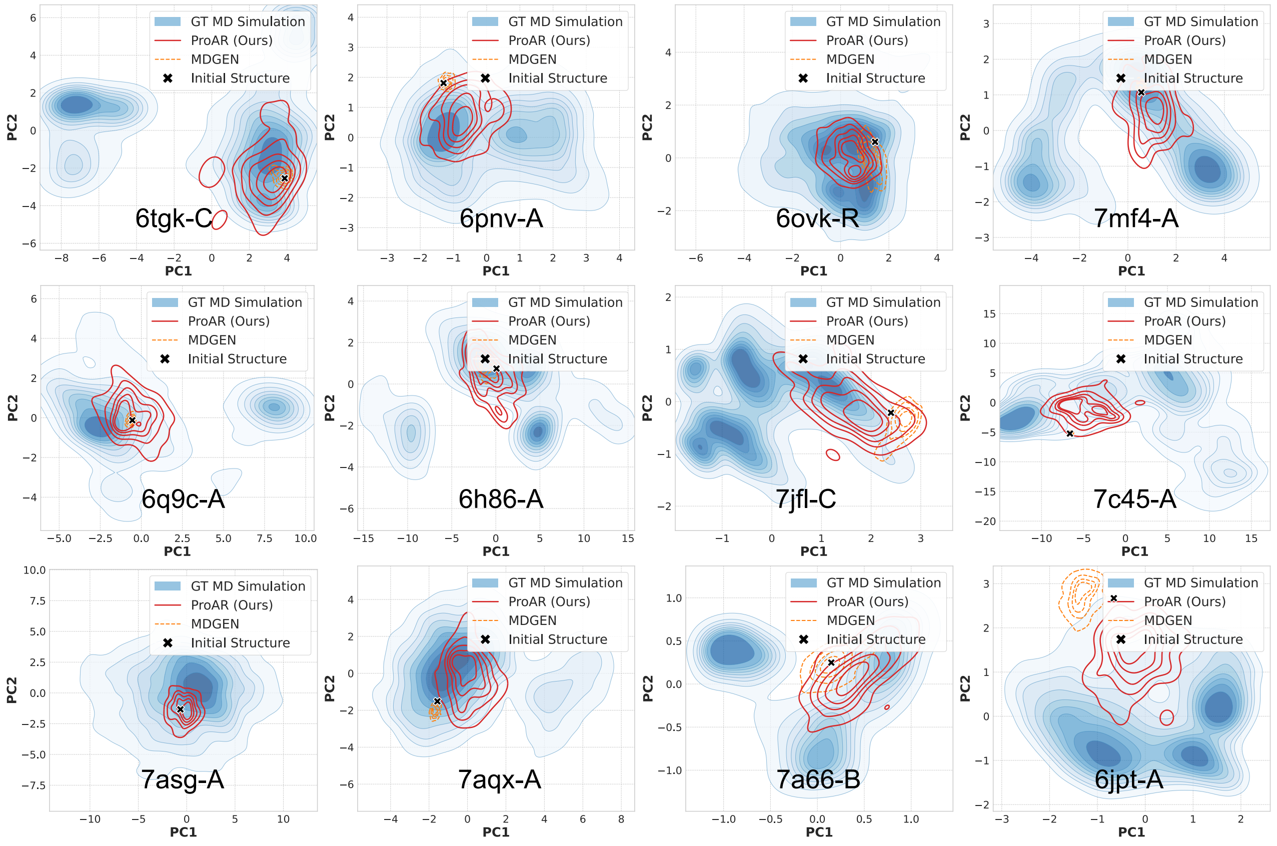


Figure 1: Visualization of the generated 100 ns trajectories for 12 randomly selected cases. The blue background represents the density of the ground-truth conformational distribution obtained from MD simulations. ProAR demonstrates improved recovery of conformational states, while MDGEN remains near the initial structure and deviates from the true distribution.

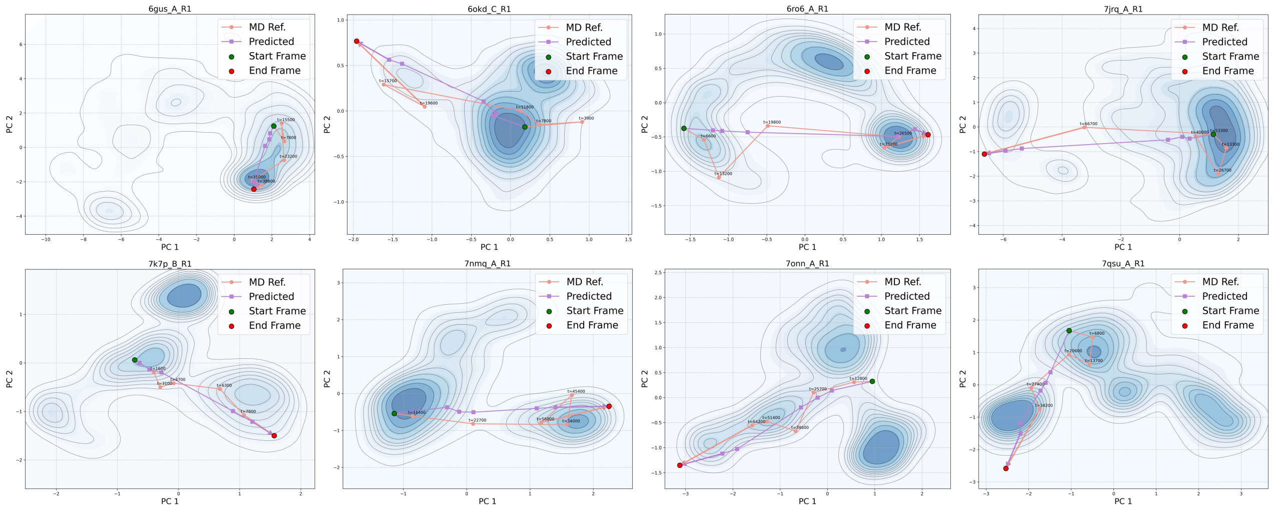


Figure 2: Interpolation results. The interpolator generates smooth pathways between the start and end frames, capturing the dynamics observed in the MD reference. The start frame corresponds to the first frame of the MD simulation, while the end frame is the conformation with the highest RMSD relative to the first frame. Although the interpolator was trained only on short segments with $h = 6$ and a 400 ps interval, it can generate transition pathways over much longer timescales. This demonstrates its ability to model transitions between arbitrary conformational states without training at multiple temporal resolutions.
